# When good guides go bad: empirical evaluation of all unique Cas9 protospacers in *E. coli* reveal widespread functionality and rules for gRNA design

**DOI:** 10.1101/2025.05.23.651106

**Authors:** Elise K. Phillips, Riley Harrison, S’Khaja Charles, Dawn Klingeman, Timothy D. Wiser, Carrie A. Eckert, William G. Alexander

**Author notes:** This manuscript has been authored by UT-Battelle, LLC under Contract No. DE-AC05-00OR22725 with the U.S. Department of Energy. The United States Government retains and the publisher, by accepting the article for publication, acknowledges that the United States Government retains a non-exclusive, paid-up, irrevocable, world-wide license to publish or reproduce the published form of this manuscript, or allow others to do so, for United States Government purposes. The Department of Energy will provide public access to these results of federally sponsored research in accordance with the DOE Public Access Plan (http://energy.gov/downloads/doe-public-access-plan).

## Abstract

The Cas9 nuclease has become central to modern methods and technologies in synthetic biology, largely due to the ease in which it can be targeted to specific DNA loci via guide RNAs (gRNAs). Reports vary widely on the actual specificity of this targeting, with some studies observing 60% of gRNAs possessing no activity against the genome, yet an assumption that inactive gRNAs are rare persists in the *E. coli* community. To resolve these contradictions, we evaluated the activity of nearly 500,000 unique gRNAs in the *E. coli* K12 MG1655 genome. We show that the overwhelming majority (at least 93%) of unique gRNAs are functional while only 0.3% are nonfunctional.These nonfunctional gRNAs exhibit strong spacer self-interaction, which can be either excluded using a simple design rule or “repaired” during library design. Finally, this work provides the greater microbial synthetic biology community both a set of nearly half a million *E. coli* gRNAs that have been empirically evaluated *in vivo* as well as a thoroughly evaluated experimental procedure, complete with appropriate controls for Cas9 activity, for conducting Cas9 assays in *E. coli* specifically and bacteria more generally.

**GRAPHICAL ABSTRACT:** 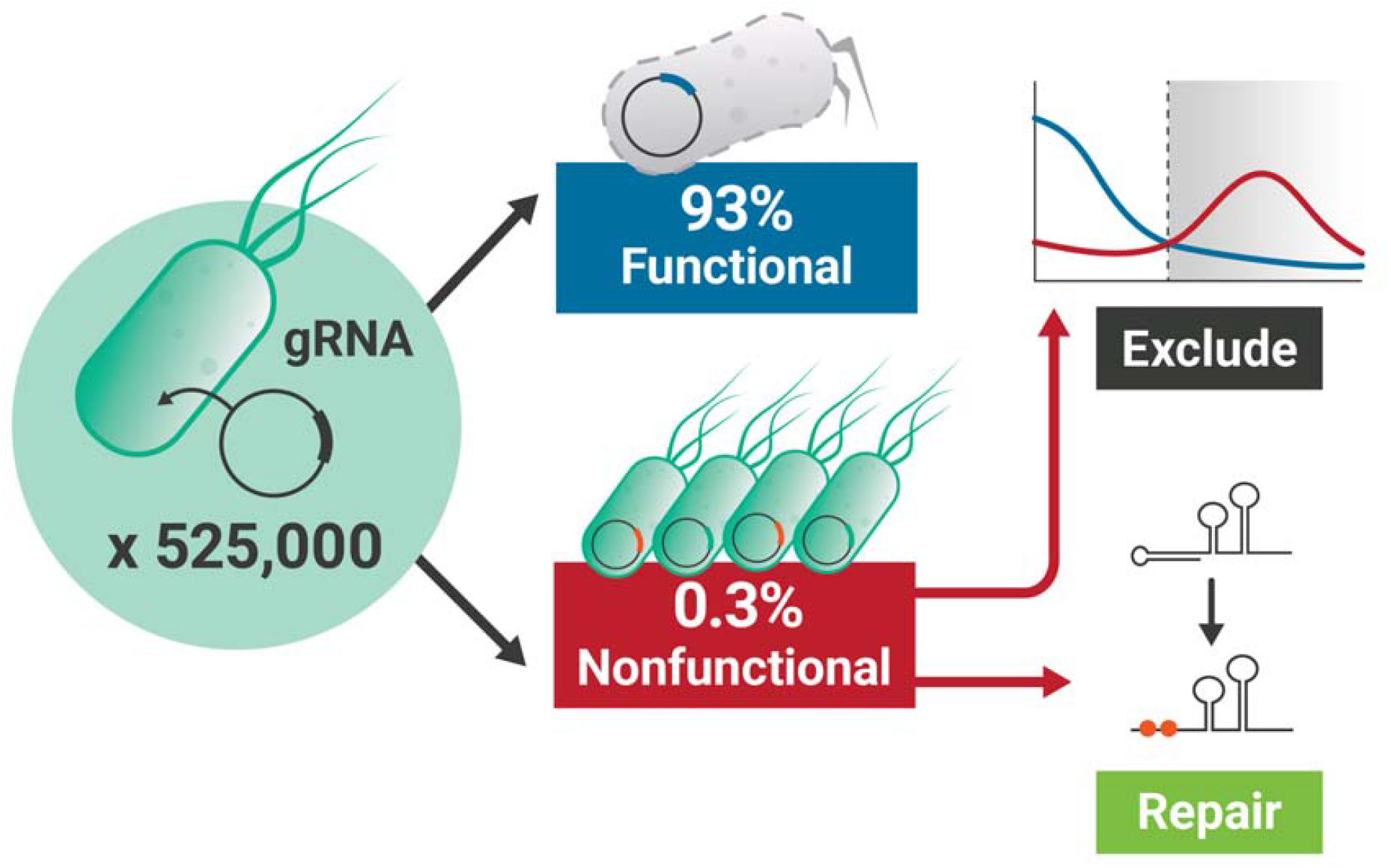

## INTRODUCTION

A ubiquitous class of genome editing techniques use endonucleases to achieve precise, arbitrary, and targeted genetic changes^1–3^. While nucleases of various types have been used in genome editing in the past, the current gold standard for targeted and precise genome editing uses RNA-directed endonucleases derived from bacterial CRISPR systems, with the Cas9 protein from *Streptococcus pyogenes* being the most well-known and widely used^4,5^. These CRISPR-associated endonucleases are directed to their targets by the binding of an associated RNA molecule to a complementary target DNA molecule^6,7^; the DNA molecule is then cut via the action of the two nuclease domains present in Cas9. For Cas9 in its native context, two RNA molecules are needed: a CRISPR RNA (crRNA) and a trans-activating CRISPR RNA (tracrRNA)^8^. However, during its development as a genome editor, the Cas9 crRNA and tracrRNA were fused together with a tetraloop to form a synthetic guide RNA (sgRNA or gRNA), which simplified heterologous use of the system^4,5^. Hereafter we refer to the RNA molecules that bind to Cas nucleases to direct their targeting generally as guide RNAs or gRNAs.

Cas9 gRNAs are 96 bases in length and contain two functional regions: the first 20 nucleotides are called the spacer, and the remaining 76 nucleotides comprise the Cas9-interaction region called the scaffold^9^. The spacer itself contains two regions: the first six bases are referred to as the Protospacer Adjacent Motif (PAM) Distal region (or merely “distal” for short), and the 14 nucleotides on the 3’ end of the spacer are referred to as the PAM Proximal region (more frequently referred to as the “seed”)^10^. Mismatches in the seed are poorly tolerated by Cas9, resulting in reduced or abolished nuclease activity, while mutations in the six distal bases can confer little to no reduction in activity^11^. Additionally, the Cas9-gRNA complex will preferentially bind and act on spacer-complementary DNA that is adjacent to a short sequence specific to that nuclease. This sequence comprises the aforementioned PAM, and for *S. pyogenes* Cas9, this sequence (NGG) must be directly adjacent to the last base of the protospacer (the locus on a genome with sequence identity to a spacer that is adjacent to an NGG motif)^12^. However, RNA-directed nucleases frequently recognize slightly altered PAM sequences, or “non-canonical” PAMs, which may serve as a mechanism to defend against selfish genetic elements evolving to eliminate PAM recognition sites from their genome. *S. pyogenes* Cas9 recognizes NAG and NGA as non-canonical PAM sequences and bind to the associated protospacer with reduced, though not eliminated, effectiveness^13,14^.

Despite the wide use of this nuclease as a genome editor for over a decade, contradiction and confusion exists in the literature and the community regarding how to choose protospacers during the design of gRNAs^15^, even in bacterial systems. For instance, up to 60% of tested gRNAs in *E. coli* have been reported as non-functional *in vivo*^16^, but within the *E. coli* community, the “lab lore” (the scientific information passed on culturally within research communities, usually verbally, and rarely published) asserts that most gRNAs just work, with little troubleshooting required. Numerous computational models have been developed to predict target suitability, but these models do not generally apply well outside of the specific contexts in which they were developed^17^.

The obvious approach to reconcile these disagreements is to test the functionality of gRNAs using a genome-scale library consisting of a sizeable proportion (or the entirety) of possible targets in a genome. The activity of members of this library can be empirically evaluated for Cas9 activity via depletion, as a functional gRNA co-expressed with Cas9 is lethal in organisms with little to no non-homologous end joining (NHEJ) capability^18,19^. Despite this utility, very few genome-scale libraries of this kind have been deployed in *E. coli,* with the largest being a 120,000-member library we previously designed and tested^17^. Despite the large relative size of this library, we still could not determine precise criteria for discriminating poorly performing protospacers from highly functional ones. Here, we designed and tested the remaining unique protospacers in the *E. coli* genome. In combination with the previously reported library, we synthesized a total of 524,596 unique protospacers, resulting in comprehensive Cas9 functionality data for 463,189, or 88%, of the total unique protospacers present in the genome. We found that almost all of the unique spacers tested (at least 93%) are highly active in *E. coli*, with only 0.3% of spacers confirmed as being inactive. We also demonstrate the role played by spacer self-interaction in gRNA inactivation and provide a method for potential gRNA reactivation to target specific loci. Finally, we develop a simple design rule to avoid the bulk of inactive gRNAs that we anticipate will be applicable to many microbial systems.

## MATERIAL AND METHODS

### General Information

All strains, primers, and plasmids used in this work are found in Supplementary Table 1. All *E. coli* strains grown in the course of this work were cultured on Lysogeny Broth (LB) – Miller Formula (5 g/L yeast extract, 10 g/L tryptone, 5 g/L sodium chloride)^20^. Agar was added to 1.5% to form solid media. Kanamycin and carbenicillin working concentrations used were 50 and 100 ug/mL, respectively, except where noted. All PCRs were performed with 2x Q5 Hot Start Master Mix (NEB, Ipswich, MA) in a C1000 Thermal Cycler from Bio-Rad (Hercules, CA).

### Design and Cloning of Oligonucleotides for Genome-wide Library Production

Three separate SurePrint OLS oligonucleotide pools were ordered from Agilent (Santa Clara, CA) to cover all unique protospacers: one containing 120,000, and two containing 202,296, resulting in three separate tubes of lyophilized single-stranded DNA. Also included in each design was the J23119 Anderson series promoter (https://parts.igem.org/Promoters/Catalog/Anderson) 5’ of the spacer and the first 35 bases of the scaffold directly 3’ to the spacer. Each oligonucleotide pool was dissolved by adding 200 µL TE buffer and incubated at 55 C for 10 minutes, then vortexing until no visible solids remained. For production of library plasmids, 20 µL of each oligonucleotide pool was added to a PCR tube along with 2.5 µL oWGA139 (which binds to the 3’ end of the scaffold), and 25 µL 2x Q5 Master Mix. The reaction was placed in a thermal cycler and the following program was run: 98 °C 1 minute, 57 °C 10 s, 72 °C 5 min, 4 °C hold, 98 °C 1 minute, [98 °C 5 sec, 65 °C 15 sec, 72 °C 15 sec] x 35, 72 °C 2 mins, 12 °C hold. During the 4 °C hold, 2.5 µL of oWGA140 (which binds to the 5’ end of the promoter) was added, the reaction pipetted to mix, then removed from the cycler and placed on ice. The program was then resumed, and the reaction was placed back into the cycler once the block temp had reached 72 °C to minimize mispriming (Figure 1A). The resultant PCR product was separated on a 3% agarose gel, the product band excised, and the DNA extracted for cloning using the Monarch Gel Extraction Kit (NEB). Note that at this step, the two larger library PCR reactions were combined equimolarly before gel purification. The gel-extracted, double-stranded library insert DNA was finally drop dialyzed for 15 minutes using 0.025 nm nitrocellulose filter from Nalgene to remove trace contaminants from gel extraction then was quantified with a Quantus fluorometer (Figure 1B).

**Figure 1.**
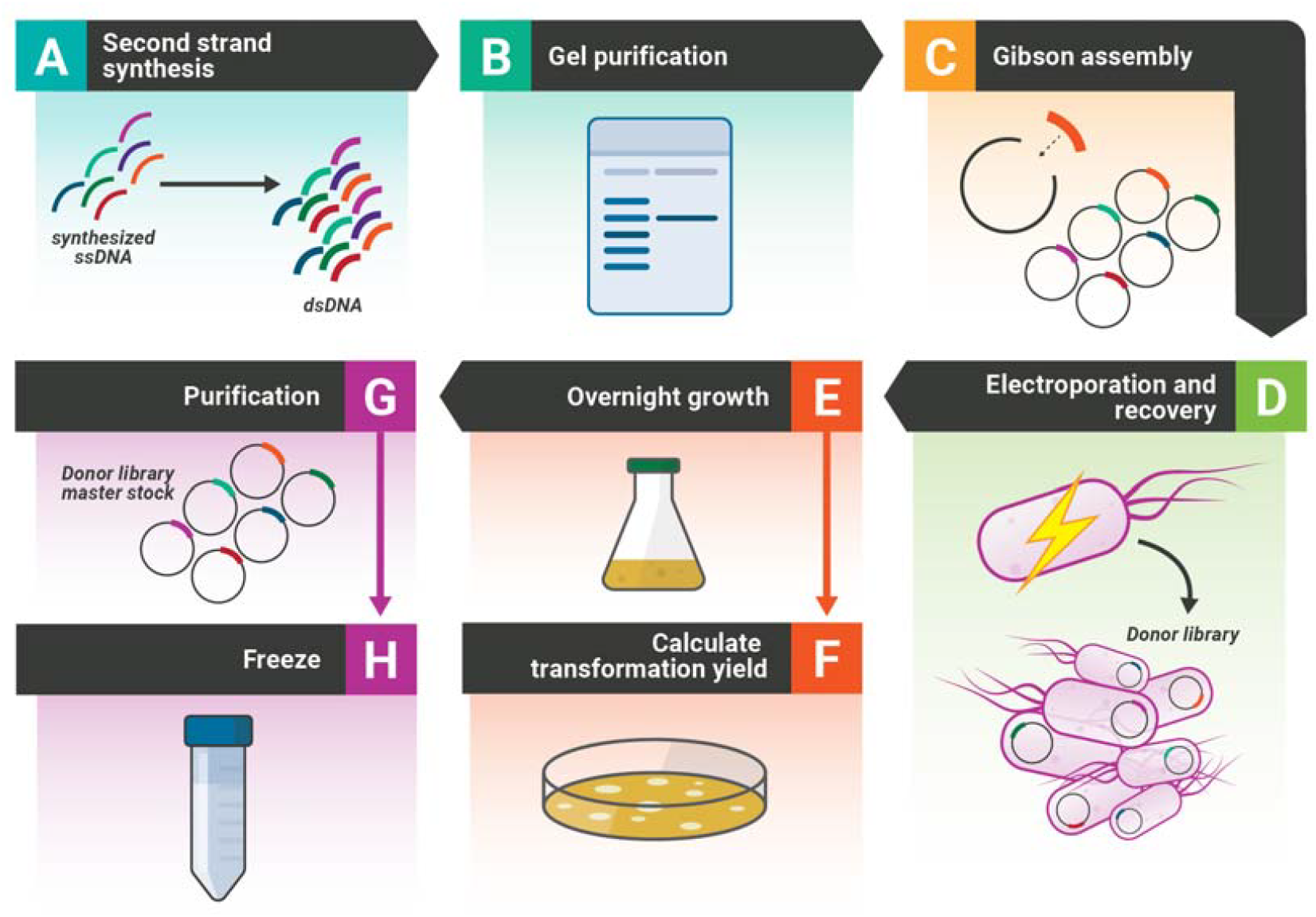
Production of donor libraries. Oligonucleotides are received as lyophilized ssDNA, which is dissolved in TE buffer. A) To begin, ssDNA oligonucleotides are converted to dsDNA then amplified via PCR with primers containing overhangs for NEBuilder activity. B) The PCR product is gel purified on 3% agarose, then the band is excised and purified. C) The purified PCR product is cloned into a library vector backbone using NEBuilder, then D) transformed into library competent *E. coli* cells with no *cas9* gene present. E) After recovery, the transformations are transferred into antibiotic containing media to select for the library plasmid backbone and left to grow overnight. F) A sample of the recovery is diluted and plated to ascertain transformation yield, which require an overnight growth as well. G) If sufficient transformation yields were obtained (∼100x or greater the size of the library), the overnight culture of donor library is both (G) extracted to produce the Donor library master stock solution and H) stored in frozen aliquots.

The library plasmid backbone was produced by amplifying pSS9-RNA with oWGA137 and oWGA138 in a Q5 reaction. After completion, 3 µL DpnI (NEB) was added to the PCR reaction, and it was incubated overnight at 37 °C. The following day, the reaction was separated and on a 1% agarose gel, and the product band was excised and gel purified. The resulting DNA was drop dialyzed on water for 15 minutes then quantified by fluorometry.

50 ng of library plasmid backbone and 8.8 ng of library insert DNA (representing a molar ratio of 5:1 insert:backbone) were combined, brought to 5 µL with water, and mixed with 5 µL 2x NEBuilder HiFi Cloning Reagent (NEB) for each individual assembly reaction; two or four individual reactions were performed for the smaller and larger sublibraries, respectively (Figure 1C). The reactions were incubated per manufacturer’s instructions, then drop dialyzed for 15 minutes on water and combined in a microcentrifuge tube on ice.

Lucigen E. cloni competent cells were purchased to produce the smaller sublibrary by transforming six 25 µL aliquots of cells with 1 µL of dialyzed assembly reaction. For the larger sublibrary, 330 mL LB was inoculated with overnight E. cloni culture to an OD_600_ of 0.1 and cultured until that value reached 0.8. This culture was split into six 50-mL conical vials, and competent cells were produced by washing with cold 10% glycerol as previously described^22^. The concentration of competent cells was standardized to 50 OD, and 50 µL of this cell suspension was electroporated with no added DNA in a 1 mm electrocuvette (Bulldog Bio, Portsmouth, NH). The electroporation was conducted with a Bio-Rad Xcel Mini Pulser using the “Ec1” program, and a time constant of 5.6 ms or greater for this electroporation was required to proceed with the transformations of the library assembly reaction (Figure 1D).

Immediately after delivering the electrical pulse, 950 µL of Super Optimal Broth (SOB^23^) was added directly into the electrocuvette. After all transformations had been performed, the resulting recovery culture was combined into a 15-mL culture tube, its volume was estimated with a serological pipette, and it was then cultured shaking at 37 °C for an hour. After recovery, 10 µL of the recovery was collected into a 1.5 mL microcentrifuge tube, then the recovery was then transferred in its entirety to a 250-mL baffled culture flask containing LB carbenicillin media, which was incubated overnight shaking at 37 °C (Figure 1E). Finally, the 10 uL sample of recovery culture was serially diluted to 10^−4^, and 10 or 100 µL of this dilution was spread on LB carbenicillin agar plates in triplicate and incubated at 37 °C.

The next day, colonies on the plated serial dilution plates were counted, and using the total volume of recovery culture transferred into the selective culture the total yield of transformants were calculated (Figure 1F). This total transformant yield is then compared to the number of designs in that sublibrary to determine depth of coverage. The library coverage is considered sufficiently deep if this yield is 100x or greater than the number of protospacers in the sublibrary being cloned (e.g. at least 12 million total transformants in the smaller sublibrary). If the library passed this quality control step, 50 mL of LB containing 200 ug/mL carbenicillin was added to the overnight culture, and it was incubated shaking at 37 °C for an additional hour. Afterwards, the culture was split into two half volumes, and both were centrifuged as before to harvest cells. One half was used with the Zymo Research Plasmid Midiprep Kit as instructed by the manufacturer (Figure 1G), while the other’s cell pellet was suspended in 21 mL LB plus 9 mL 50% glycerol and split equally among three 15-mL conical vials. These three aliquots were flash frozen with liquid nitrogen for long term library storage, and the resulting plasmid miniprep is termed a donor library (Figure 1H).

### Transformation of Genome-wide Library in *E. coli* MG1655 Host Strain

We standardized the donor plasmid library to a working concentration of 20 ng/µL to improve reproducibility. “Spike-in” control plasmids (pWGA128 and pWGA130) that had been extracted prior from a clonal source were added at a concentration to match that of individual designs as determined by dividing the library mass used (100 ng) by the number of protospacers in the library. The mass of control plasmids used was ∼0.83 pg/100 ng library plasmid for the smaller sublibrary and ∼0.25 pg/100 ng library plasmid for the larger sublibrary. The spike-in controls have been validated for nonfunctionality and functionality, respectively, and serve as internal standards for the functionality of gRNAs tested in the libraries.

The *E. coli* K12 MG1655 host (hereafter MG1655) used contains the pX2-Cas9 plasmid, which drives Cas9 expression with the arabinose-inducible promoter P_araBAD_. For transformation, electrocompetent cells were prepared as described for the donor library, except that MG1655 was grown on LB +50 ug/mL kanamycin +0.2% arabinose. Ten host electroporations were performed as before for the smaller sublibrary, and thirty were performed as before for the larger (Figure 2A). After combining all transformations for recovery, the wash control plasmid pWGA129 is added to a concentration of 1 ng/mL of recovery volume; this plasmid is used to determine the contribution of extracellular DNA, if any, to the data collected on these libraries (Figure 2B). After recovery at 37 °C for two hours, the entirety of the culture is transferred to 50 mL LB + kanamycin +carbenicillin in a 250 mL baffled flask then shaken at 37 °C overnight (Figure 2C). The next day, 50 mL of cells are harvested, washed twice with 50 mL DNase Wash Buffer (10 mM Tris, 2.5 mM MgCl_2_, 0.5 mM CaCl_2_, pH 7.5), then resuspended in 995 µL of the same buffer and transferred to a microcentrifuge tube. 5 µL of DNaseI (NEB) was added, and the tube was incubated at 37 °C for 15 minutes (Figure 1D). Afterwards the cells were harvested by centrifugation, the supernatant was removed, and DNA was extracted from the cell pellet using the Zymo Research Midiprep Kit (Figure 2E) for amplicon sequencing by Illumina (Figure 2F).

**Figure 2.**
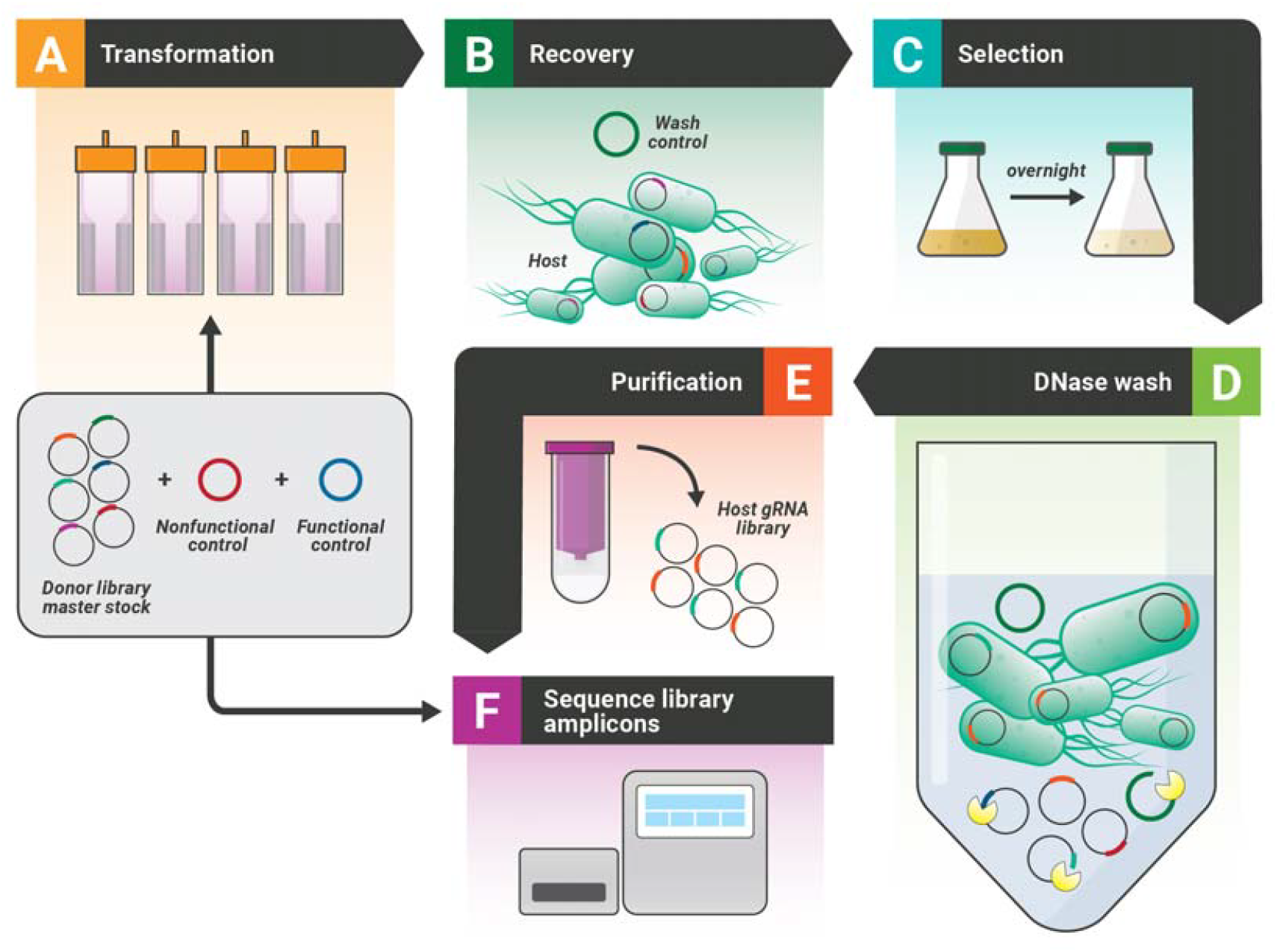
Production and analysis of a host library. Donor library master stock solution is “spiked” with functional and nonfunctional gRNA control plasmids, then A) 3 to 30 transformations of the host (induced to express Cas9) are performed. B) All transformations are combined and recovered in a single culture tube, and the wash control plasmid is added at a concentration of 1 ng per uL of recovery culture. C) After recovery, the entirety of the recovery culture is added to LB +kanamycin +carbinicillin and incubated overnight. D) The next day, a sample of the overnight culture is harvested, the cell pellet washed twice with DNase Wash Buffer, then suspended in the same and treated with 5 uL DNase I to degrade any DNA not contained within a living cell. The cleaned cell pellet is E) immediately extracted for plasmids, then F) both the Host library and the Donor library containing controls are sequenced on an Illumina platform.

### Phased PCR Amplicon Sequencing Analysis of Genome-wide Libraries

Amplicon sequencing is challenging for Illumina instruments, as low library diversity causes poor sequencing cluster differentiation, leading to a loss in individual cluster resolution and read quality. gRNA libraries are considered to be low diversity (and thus problematic to sequence) because the scaffold and promoter are identical across all members. As Illumina sequencing of amplicons is currently the only method available to assess these libraries, we developed phased PCR amplicon sequencing to address these technical issues. We constructed forward and reverse primer sets that possess between zero and four random bases added on the 5’ end. These random bases are sequenced first by the Illumina instrument, meaning that instead of all clusters registering the same nucleotide, an assortment of all four bases will be observed. For additional diversification of the library, we prepared the samples with the NEB Illumina Ultra II kit, which attaches hairpin adapters in random orientation relative to the start and end of the cassette and included 20% PhiX control DNA. Each sequencing library was sequenced in duplicate as a technical replicate to determine measurement errors. Each sublibrary replicate was sequenced to a minimum read depth of 100x the number of designs in that sublibrary. To obtain this read depth the smaller sublibrary was sequenced with a single MiSeq, while the larger sublibrary was sequenced using both lanes of a NextSeq flowcell.

For each donor and host library replicate, paired-end reads were merged using USEARCH^24^ and the -fastq_merge command, and the number of merged pairs was recorded (Figure 3A). The merged reads were then searched against the index of gRNA designs for that library with the USEARCH -usearch_global command, setting *id* to 0.9 and target_cov to 0.9. Both strands were searched, and the single top hit for every merged read was reported. The -userout custom output was configured to report the read name queried, the name of the target protospacer match, and the proportional identity of the match. This custom output file was then used as input for a bespoke Python script that compiled the raw data into a count of full length, perfect matches (matches that are identical to a protospacer design over the entire length of the protospacer) for each design in the library (Figure 3B). The script also takes the number of merged reads as an argument to calculate the abundance of each design (number of full length perfect matches to a design/total merged pairs); if directed by the user, the script will replace a count of 0 with 0.5 during the abundance calculation, which is used on host library data as a workaround for statistics errors arising from zero values^25^.

**Figure 3.**
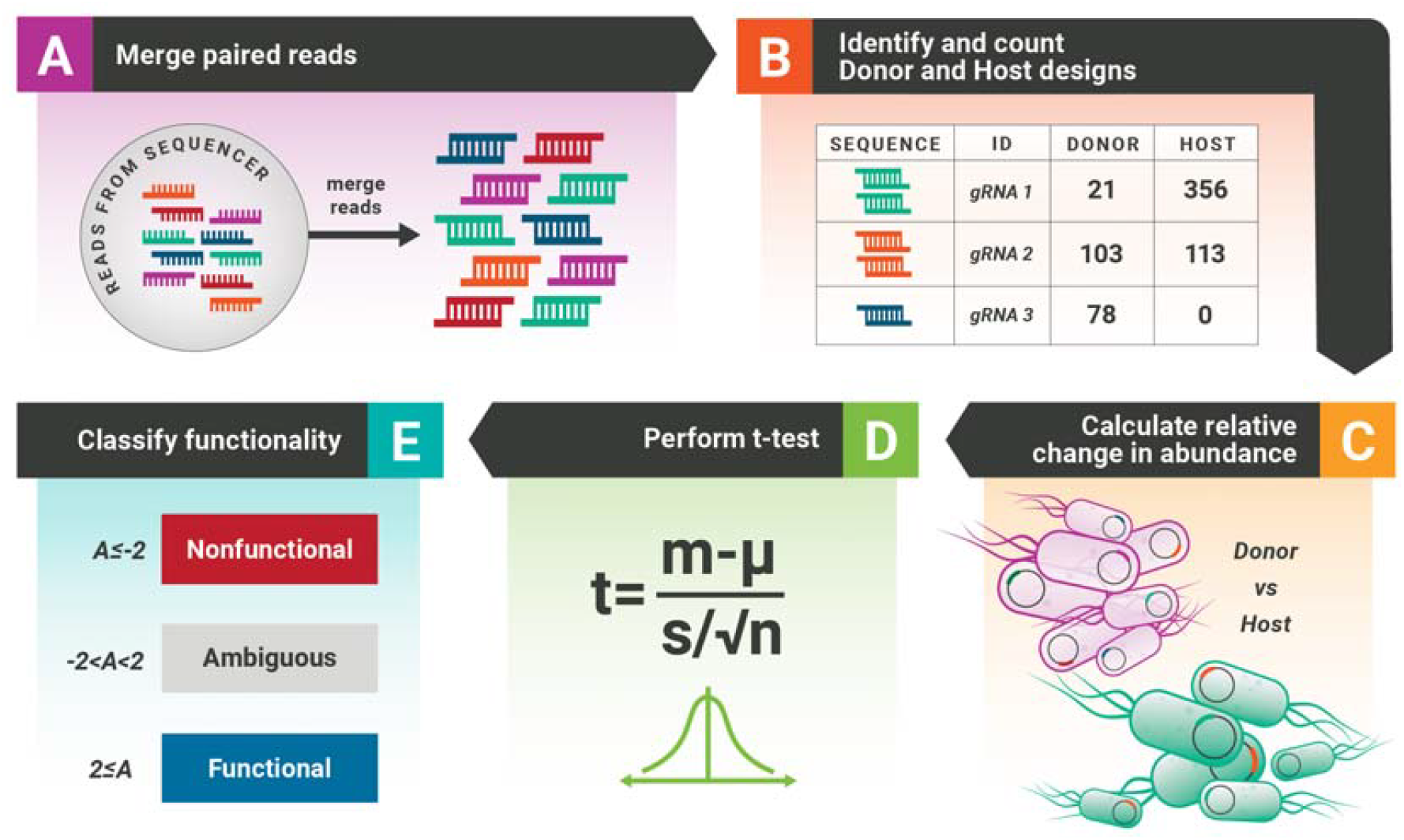
Host library evaluation scheme. A) Paired end Illumina reads are merged then B) identified with USEARCH. A custom script parses the output of USEARCH to produce a CSV file containing raw Illumina read counts and C) calculated abundance for each design in the library. D) These data are collated, statistical tests for significant changes in abundances are run, and E) each design is classified by functionality based on various metrics.

The abundance values for all designs in all replicates of donor and host were compiled into a single spreadsheet for analysis. Per design, the average host abundance was divided by the average donor abundance, then the -log2 of this change in abundance was calculated (Figure 3C). Also, a Student’s T-test was performed (a=0.05) on the donor vs. the host abundance values per design to determine statistical likelihood of the differences observed between donor and host being due to chance, and designs with P>0.05 were labelled as “Statistically Insignificant” and excluded from further analysis (Figure 3D). Also excluded from analysis were any designs with average donor library read counts of less than 20, as small changes in read depths in these designs would result in oversized changes in apparent abundance; this value was chosen largely arbitrarily, though similar values have been used in prior works (Bassalo et al., 2018; Garst et al., 2017). gRNAs were labelled as functional if they exhibited statistically significant change in abundance and had an A-score >= 2. A nonfunctional label was instead applied if a gRNA exhibited significant change and had an A-score <= -2 (Figure 3E).

### Production and Analysis of the Confirmation and Nonfunctional Libraries

The Confirmation Library was designed by randomly choosing 200 functional, 200 nonfunctional, and 200 statistically insignificant gRNA spacers from the Genome-wide library. Library oligonucleotides were ordered from Genscript (Piscataway, NJ) and used to produce a donor library using the same method as the Genome-wide library, except that only three transformations of competent cells were used. Working donor stock production and host cell transformation were performed as described with the Genome-wide library, except again only three transformations were conducted. Sequencing was prepared as before and conducted using a MiSeq. Analysis was performed as before with the Genome-wide Library.

The Nonfunctional Library was made and analysed in a manner identical to the Confirmation Library, with the following alterations: 1) all 2,365 putative nonfunctional spacers were ordered for synthesis, 2) three technical replicates of donor and host were sequenced rather than two, and 3) a new set of phased PCR amplicon sequencing primers were used. These new primers were designed to a) avoid issues with priming on regions of interest and b) to produce an amplicon containing the entirety of the promoter, spacer, and scaffold of a gRNA. The lattermost two alterations were done in efforts in improve the accuracy of functionality calls for this library.

### Spacer self-interaction prediction

ViennaRNA^26^ with default settings was used to predicted minimum free energy (MFE) and secondary structure for all spacers demonstrating statistically significant differences. These data were then binned by MFE values and plotted.

## RESULTS

### Rationale of Approach

gRNA functionality can be evaluated in a massively parallel fashion through the production of gRNA libraries via oligonucleotide synthesis, which produces rationally designed, custom DNA molecules. When cloned into a plasmid backbone (Figure 1) and transformed into a target bacterium expressing Cas9 (Figure 2), functioning gRNAs will mediate Cas9 activity at the corresponding protospacer, resulting in cell death unless repaired. Thus, in organisms where NHEJ is absent, functional gRNA designs will decrease their proportion in a population over time; reciprocally, gRNAs that do not trigger Cas9-mediated cell death will increase their relative proportions (or abundances) as those designs behave like an empty vector. Using high-throughput sequencing, we can track the changing abundances for all designs in a library, and by comparing those changes to those of known control designs spiked into the library we can infer functionality of each tested design (Figure 3). In short, functional gRNAs will disappear from our sequencing dataset while nonfunctional ones will be retained.

### Production of a Comprehensive Protospacer Database for *E. coli*

A bespoke Python script was written to find all unique gRNA targets within a genome. The input for this script is a FASTA formatted file which contains the genomic sequence to search for protospacers. The script examines every adjacent pair of nucleotides for identity, and when canonical (GG for a “forward” directed PAM, CC for a “reverse”) or noncanonical (AG, GA, CT, TC) PAM dinucleotides are found, the script captures the adjoining protospacer seed sequence (hereto referred as the protoseed). Protoseeds from noncanonical PAMs are stored in an exclusion set, while captured canonical protoseeds are compared the noncanonical exclusion set, a set of presumed unique protoseeds, and a set of nonunique protoseeds. If no exact match is found in the presumed unique set and the queried protoseed is not otherwise excluded, it is added to the presumed unique set. If a match is found, then the matching sequence is removed from the presumed unique set and recorded in the nonunique exclusion list. These lists persist when the script moves between replicons in the genome, ensuring that only unique protoseeds are present within the dataset. Once all unique protoseeds have been found, the script compiles information about the location, strand and sequence of the corresponding protospacer for each protoseed into a pandas^21^ DataFrame, which is then exported as CSV and a FASTA file. For this work, 524,596 protospacer sequences were found in the *E. coli* genome.

### Massively parallel depletion/enrichment analysis elucidates gRNA activity

Towards the development of comprehensive and reliable data on gRNA functionality *in vivo* in *E. coli*, we designed a library of gRNAs to target all potential unique spacers present in the *E. coli* genome. A total of 524,596 unique protospacers were identified using the above-described bespoke Python script and this set was synthesized as three separate oligonucleotide sublibraries. Multiple sublibraries were synthesized due to the maximum library size available from our oligonucleotide vendor (Agilent, Santa Clara, CA) being 244,000 designs. A smaller sublibrary of 120,000 designs was initially tested in a previously reported study^17^, and the remaining designs were produced as two 202,298-member pools. Results reported here are a summation of the results from all three libraries.

The gRNA libraries were combined then cloned into *E. coli* without *cas9*, termed the donor library (Figure 1). This library was comprised of a total of 471,530 designs, or 90% of unique protospacers designed, which passed the cutoff of at least 20 reads in both sequencing replicates. After the gRNA libraries were transformed into the host strain of *E. coli* (MG1655 with an inducible *cas9* plasmid (Figure 2)), 463,189 spacers exhibited a statistically significant change in abundance from the donor library (α=0.05). Due to size constraints, a subset of these data are shown in Table 1. These changes in abundance were -log2-transformed to aid in comparison and visualization. With this -log2 transformation, functional gRNAs have a higher positive score, and nonfunctional gRNAs have a more negative score. Of those gRNAs with significant changes in abundance, 438,495 (93%) of the spacers were classified as functional, as they exhibited a sizeable decrease in abundance (A-score >= 2) when transformed into their host, due to genomic cutting by Cas9. Additionally, 2,344 (0.5%) of the spacers demonstrated nonfunctionality (A-score <= -2) by increasing in abundance because without significant Cas9 activity cells containing these gRNAs survive and replicate. The remaining 22,350 (6.5%) of spacers were ambiguous, exhibiting relatively small yet statistically significant differences in their abundance, with A-scores between -2 and 2 (Supplementary Data 2). Surprisingly, this data indicates that the vast majority of unique spacers in *E. coli* are highly effective and functional (Table 2).

**Table 1.**
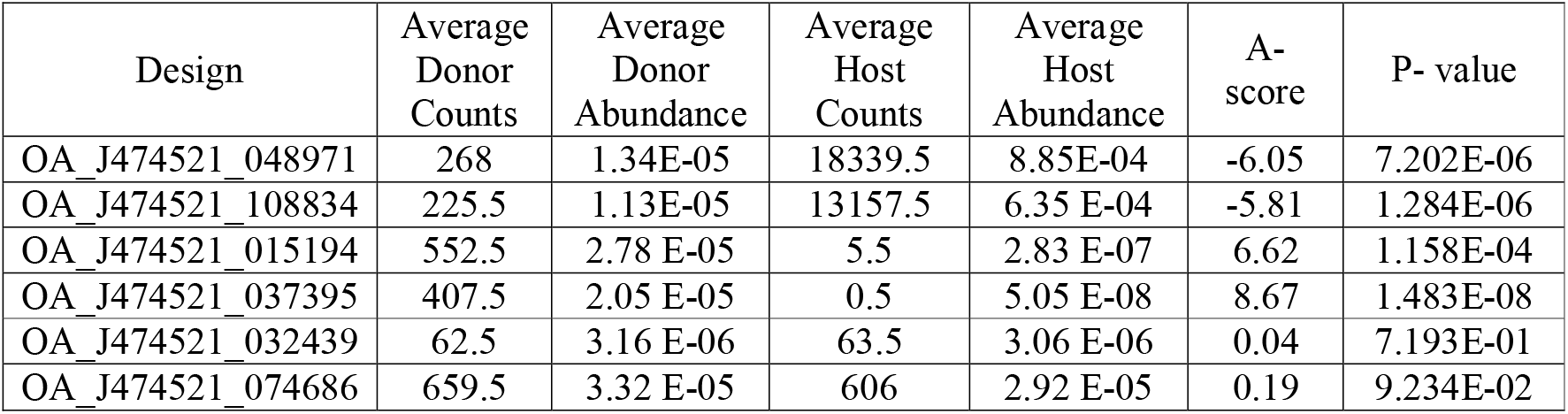
Selected excerpts from the Supplementary Data 2. For each functional category, two representative protospacer designs were manually chosen to illustrate commonly seen dynamics in large gRNA libraries.

| Design | Average Donor Counts | Average Donor Abundance | Average Host Counts | Average Host Abundance | A-score | P-value |
| --- | --- | --- | --- | --- | --- | --- |
| OA_J474521_048971 | 268 | 1.34E-05 | 18339.5 | 8.85E-04 | -6.05 | 7.202E-06 |
| OA_J474521_108834 | 225.5 | 1.13E-05 | 13157.5 | 6.35 E-04 | -5.81 | 1.284E-06 |
| OA_J474521_015194 | 552.5 | 2.78 E-05 | 5.5 | 2.83 E-07 | 6.62 | 1.158E-04 |
| OA_J474521_037395 | 407.5 | 2.05 E-05 | 0.5 | 5.05 E-08 | 8.67 | 1.483E-08 |
| OA_J474521_032439 | 62.5 | 3.16 E-06 | 63.5 | 3.06 E-06 | 0.04 | 7.193E-01 |
| OA_J474521_074686 | 659.5 | 3.32 E-05 | 606 | 2.92 E-05 | 0.19 | 9.234E-02 |

**Table 2.** Summary of gRNA functionality classification of the Genome-wide Library

| A-score | Number of gRNAs | Percent of gRNAs | Classification |
| --- | --- | --- | --- |
| $\geq 2$ | 438,495 | 93 | Functional |
| $\leq -2$ | 2,344 | 0.5 | Non-functional |
| -2 to 2 | 22,350 | 6.5 | Statistically Insignificant |

To coarsely evaluate and visualize the distribution of this activity data across the genome, we divided the *E. coli* genome into 2.5 kbp bins starting at the first base (*e.g.* 1-2500, 2501-5000, etc.). For each bin, the number of unique gRNA targets (the spacer and PAM sequence on the genome that is acted upon by the gRNA:Cas9 ribonucleoprotein complex), the number of functional gRNA targets, and the number of nonfunctional targets were totaled and plotted, along with other common genomic features, into a circos plot (Figure 4). The ratio of functional to nonfunctional gRNA targets largely remained the same across the genome, though the total number of gRNAs did vary as a function of GC content in a given region, as expected with Cas9 recognizing an NGG PAM. The sole exception to this trend is a region between 280 and 290 kbp which is enriched for both nonfunctional and total number of targets (Supplementary Note).

**Figure 4.**
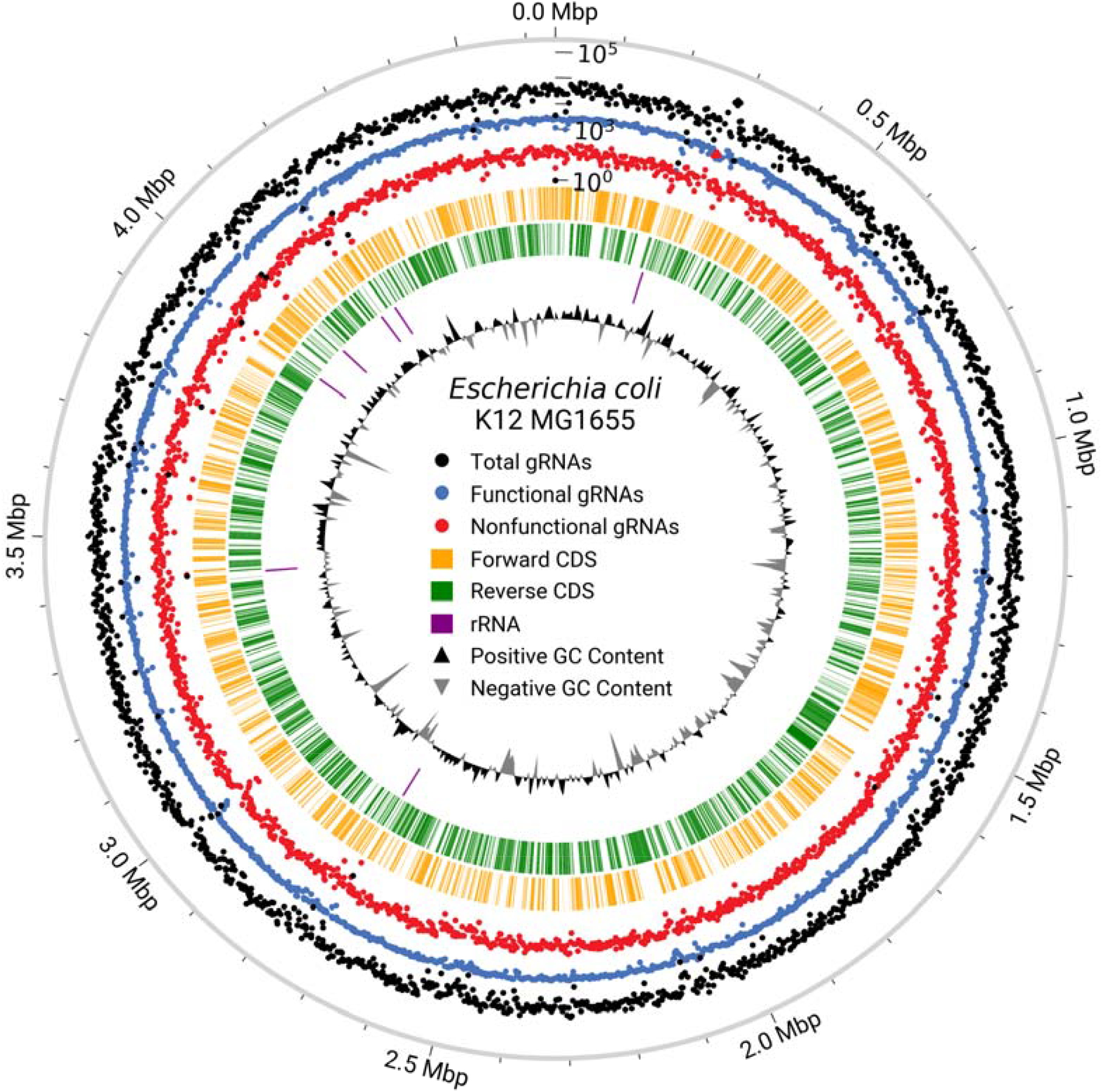
Functional and nonfunctional gRNA targets are uniform across the genome. A bespoke Python 3 script was written using the pycirclize^27^ and matplotlib^28^ libraries to generate a circos plot. The outer ring contains the genomic coordinates for the *E. coli* K12 MG1655 genome. A triple semi-log scatter plot illustrates the number of total (black dots), functional (blue dots), and nonfunctional (red dots) gRNAs present in each bin. The central dashed lines represent the location of protein-encoding (yellow and green), tRNA (pink), and rRNA (purple) genes. The internal line plot outlines the GC content of each region.

### Validating massively parallel data with smaller, more accurate pools

To validate results from the pooled library, we generated smaller library to ensure the results are repeatable and accurate. For this “Confirmation library”, we chose two hundred gRNAs each that were classified as functional or nonfunctional along with two hundred gRNAs that were statistically insignificant in their abundance changes (classified as ambiguous). These lattermost designs were included as they were somewhat paradoxical: a functional gRNA is lethal, so should exit the population while a nonfunctional gRNA increases in abundance because it behaves like an empty vector, so a design showing no significant change of abundance is contradictory. When we transformed this library into our host strain, almost all (99%) of the functional gRNAs replicated their functionality (Figure 5, Supplementary Data 3). Surprisingly, 170 designs previously classified as statistically insignificant (85%) and 150 nonfunctional designs (75%) demonstrated functionality in this library, which suggests that the cutoff value used in the genome-wide library functionality call is conservative in nature.

**Figure 5.**
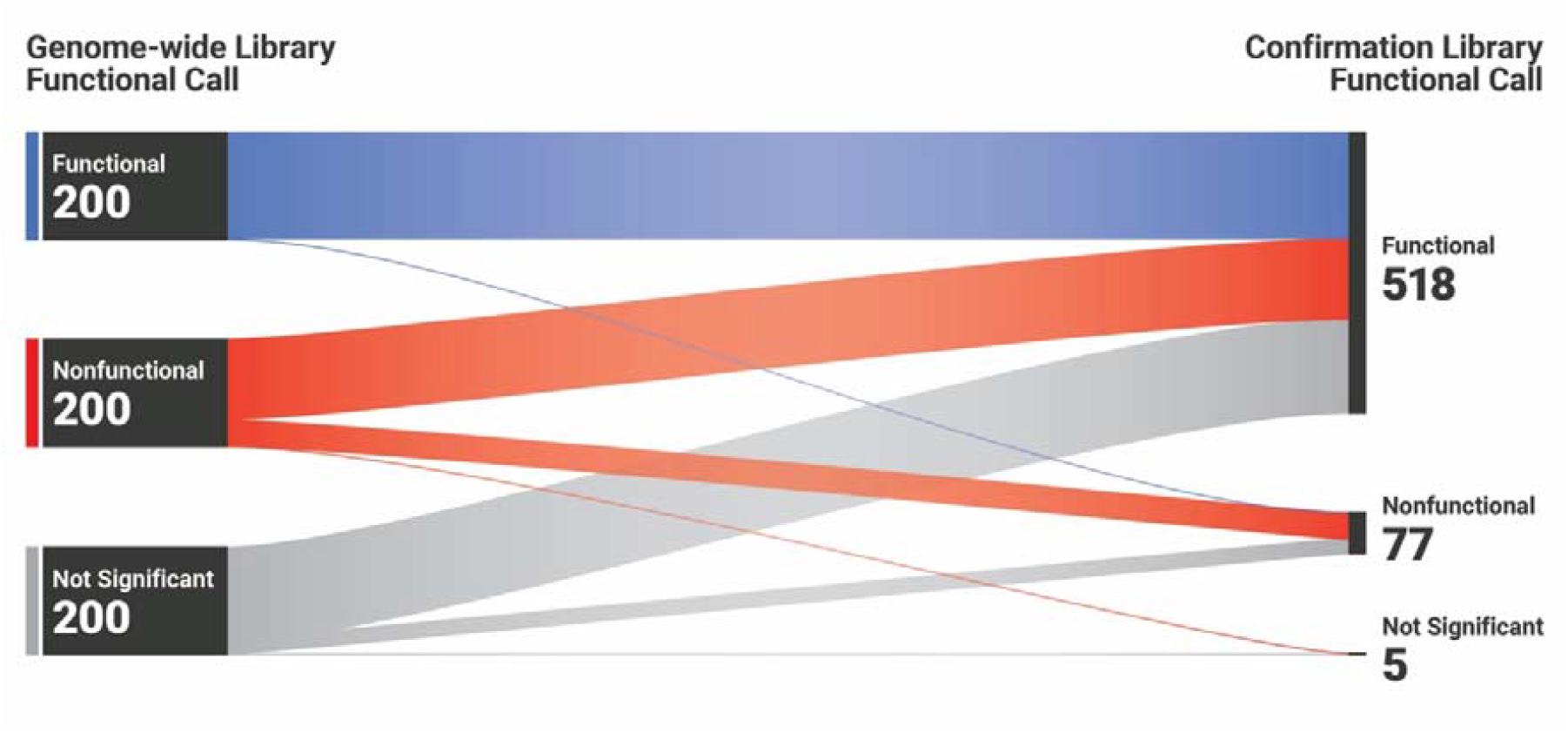
Functional calls from the genome-wide library are conservative. Randomly chosen gRNAs from the genome-wide library (200 each functional, nonfunctional, and statistically insignificant designs) were resynthesized by Genscript then cloned to produce the Confirmation library, which was then transformed into *E. coli* K12 MG1655 and analyzed. While almost all of the putative functional gRNAs demonstrated functionality in the Confirmation library, most of the putative nonfunctional and statistically insignificant designs demonstrated strong functionality in the Confirmation pool.

### Confirming poor guide activity via improved detection methods

The results from this smaller Confirmation library largely corroborate those from the genome-wide library: almost all unique gRNA targets in the *E. coli* genome result in cutting by Cas9, in contradiction to previous reports^16^. However, some of the putative nonfunctional guides repeated that behavior in this smaller library, meaning that nonfunctional gRNAs do exist in this library albeit in small quantities. As the number of inactive gRNAs were quite small (2,344) and we were interested in discovering what makes these gRNAs poor performers, we synthesized these gRNAs and cloned them to form the Nonfunctional library. During design of this library, an error in sorting data resulted in the inclusion of a small number of gRNAs that should have been filtered out due to low Donor library counts; we have decided to include these to simplify analysis, bringing the total number of Nonfunctional library designs to 2,375. We also modified our sequencing methodology for this library because we hypothesized that our sequencing method, which used amplicons primed partially on the gRNA scaffold, could be masking mutations arising within the gRNA scaffold, which would result in a misclassification of a mutated gRNA design as “inactive” when otherwise it would be excluded during analysis. We addressed this potential inaccuracy in the Nonfunctional library analysis by 1) sequencing the entirety of the gRNA cassette and 2) sequencing both Donor (from the library cloning strain, Figure 1) and Host (from the experimental strain containing *cas9,* Figure 2) libraries in triplicate rather than duplicate to further decrease the impacts of potential variance. Of the 2,375 gRNAs tested in the Nonfunctional library, 1,651 exhibited nonfunctionality under these evaluation conditions (Supplementary Data 4). These data show that only 0.3% of the total number of unique gRNA targets in the *E. coli* genome are nonfunctional for Cas9 targeting and cutting.

### Restoring gRNA activity by disrupting spacer self-interaction

One hypothesis for the nonfunctionality of these gRNAs is the presence of secondary structure in the spacer region, which has previously been shown to inhibit gRNA activity^29–31^. To test this hypothesis, RNAfold^26^ was used during the design process to predict the structure and minimum free energy (MFE) for spacers in the Nonfunctional library. We observed that most nonfunctional gRNAs had more negative predicted MFEs than the functional gRNAs. Subsequently, ten nonfunctional gRNAs were chosen to rationally mutate with increased MFE (reduced stability and secondary structure) to directly measure the impact of MFE on functionality. These designs were manually picked because they had stretches of consecutive base pairing between distal and seed regions as well as a low MFE, indicating very stable spacer structure. Thus, these gRNAs are likely to be confirmed as poorly functional, and they could hypothetically be restored to function by mutating one or two bases in the distal region of the spacer, which is less important for targeting specificity. Specifically, if one or two mutations could disrupt the secondary structure stability, the mutated gRNAs should be functional. To test this approach, ten “repaired” gRNAs were designed to test alongside their predicted nonfunctional counterparts in the Nonfunctional library (Table 2, Supplementary Data 4). The repair involved manually choosing a base most likely to destabilize the spacer via hypothetical base stacking disruption, then confirming a predicted increase in MFE via RNAfold.

All ten subject gRNAs matched our initial, manual prediction of non-functionality, and nine of the ten repaired versions of these gRNAs were restored to functionality as shown by a marked depletion after transformation into the host (Table 3, Supplementary Data 4). This suggests that if a gRNA designed to target a specific region has high likelihood of secondary structure and therefore nonfunctionality, the design could be slightly altered to become functional. This is especially true because nucleotide base stacking in very stable gRNAs is not additive, so minor changes, as little as one nucleotide, can lead to major improvements in gRNA function.

**Table 3.**
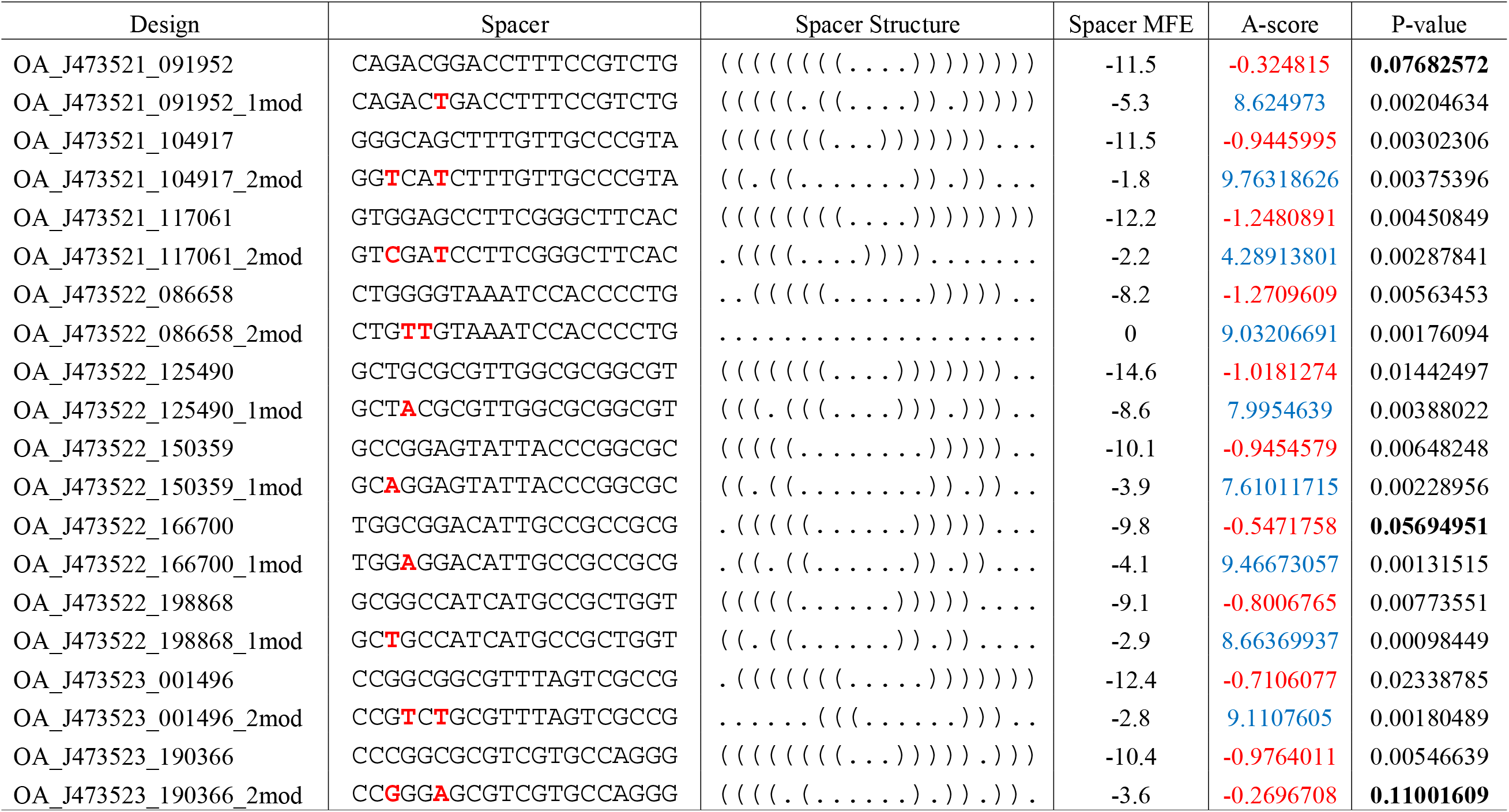
Nonfunctional gRNAs can be restored to function by disrupting spacer secondary structure. Mutations made are in red and emboldened. Functional A-scores are colored blue, while nonfunctional scores are red. Finally, statistically insignificant P-values are emboldened.

### Developing design rules for gRNA libraries

Our results validating that spacer self-interaction has major effects on the function of a gRNA suggest that the predicted MFE value can act as a proxy for a number of physical effects that cause strong self-interaction (number and identity of bases paired, base stacking effects, etc.). Thus, we wanted to determine whether any trends emerged that could be used as gRNA design rules. To compare the distribution of MFE values in all functional gRNAs to the 1,651 confirmed nonfunctional, we separated all gRNAs in both functional categories into bins based on their spacer MFE, tallied the number of spacers in each bin for each category, and then plotted them (Figure 6A). While the vast majority of functional gRNAs had an MFE value of -5 kcal/mol or greater, the MFE of the nonfunctional gRNAs peaked at -9 kcal/mol. Graphing these bins we observe very few total gRNAs with values between -7 an-6 kcal/mol. This shows that an MFE cutoff of between -6 and -7 could serve as a simple but effective cutoff during library design to avoid poorly performing guides with little loss of high-performing gRNAs (Figure 6b). -6.5 kcal/mol seems ideal, as this value would reject 61% of nonfunctional gRNAs while only losing 2.7% of functional gRNAs. Depending on specific experimental contexts and demands on library design a more or less strict cutoff could be appropriate.

**Figure 6.**
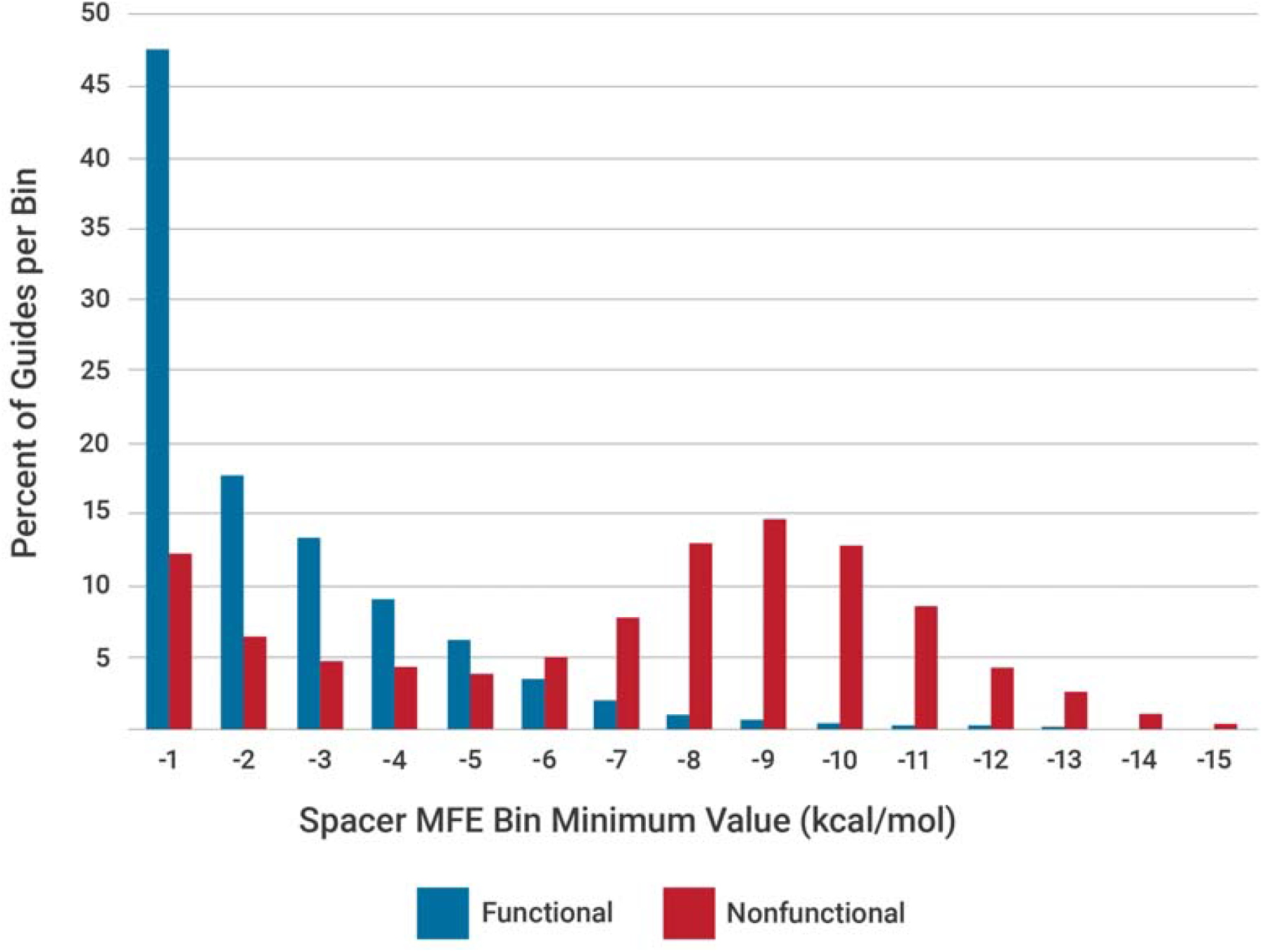
Functional gRNA spacers exhibit greater minimum free energy values than nonfunctional spacers. A. Spacer MFE values for both functional and nonfunctional gRNAs were binned, then the total for each bin was normalized to the total. The resulting percent of total gRNAs for each bin was then plotted.

## DISCUSSION

Here we detail findings from the first comprehensive screen of Cas9 gRNA functionality in the *E. coli* genome *in vivo*. We show that the majority of unique gRNAs (over 93%) are functional. While the vast majority of Cas9 targets in *E. coli* can be acted upon effectively, we also demonstrated the feasibility of restoring nonfunctional gRNAs to functionality by altering bases in the distal region and determined an empirically informed cutoff MFE value of -6.5 kcal/mol to best avoid the poorest-performing gRNA designs. Finally, we present to the synthetic biology community close to half a million gRNA sequences that have demonstrated activity *in vivo* in *E. coli*.

While many gRNA design software packages and associated rules are available for gRNA design, they are often unreliable in practice. Numerous factors likely contribute to this disfunction, though misapplication of models derived from unrelated biological systems is arguably the most common and most obvious. For example, the commonly used Doench score^32^ is one of the first attempts to quantify gRNA target suitability, but the datasets gathered were from immortalized human cell lines. Despite the very specific context in which this model was developed (eukaryotic system, heavily mutated with compromised cell cycle control and aneuploidy), it sees widespread application as a default in many popular biological CAD programs (*e.g.* Geneious, Benchling) regardless of the user’s biological context (e.g. bacteria, plants, archaea, *etc.*). One likely source for gRNA misclassifications in bacteria is the role of chromatin structure, which is likely factored into models developed for eukaryotes. As chromatin structure has been shown to impede Cas9 ribonucleoprotein access to DNA both *in vivo* and *in vitro*^31,33,34^, the model employed to generate a Doench score will likely predict features correlated with nucleosome occupancy in the *E. coli* genome spuriously. These spurious, out of context features will then mis-score a relevant gRNA. Our results also suggest why the Doench score and other inappropriate measures of gRNA suitability were able to be used for so long in *E. coli* design: if greater than 93% of the potential gRNAs made for *E. coli* are functional, the criteria used to pick functional gRNAs ceases to matter. Also, for those gRNAs flagged as poor by models, a potentially large pool of gRNAs is excluded from testing because the software predicted they would be nonfunctional, thus inflating the number of gRNAs that people believe to be bad.

In addition to the possible classification disparities from using the wrong prediction tools for gRNA design in *E. coli,* there are other potential reasons that our results here contradict previous reports detailing poor gRNA activity within *E. coli*^16^. Cas9 expression conveys a mild fitness defect, which has also been reported previously^35^ but is a familiar phenomenon for those who work frequently with Cas9 in living systems. Issues with this defect manifest when Cas9 expression is constitutive and/or too high, as cells will mutate constructs driving Cas9 expression to relieve this defect. Cells that reduce or eliminate this growth defect by mutating Cas9 constructs obtain improved fitness compared to the rest of the population, in turn causing an over-representation in the library. Further, this Cas9-mutant, if it possesses a gRNA library plasmid, will then replicate that plasmid as it continues to grow, and the gRNA produced appears to be nonfunctional since no Cas9 is expressed.

This phenomenon could explain previous reports of high levels of nonfunctional gRNAs^16^, as a Cas9^-^ mutant will behave exactly as a nonfunctional gRNA with functional Cas9. These selective pressures must be accounted for during experimental design and, if possible, worked around to prevent these “escapees” from confounding the data. Our use of arabinose-inducible Cas9 in this work allows us to control the duration and level of expression to avoid generating these escapees. Additionally, appropriate and informative controls is crucial to interpret library data correctly. The data presented here were produced using three “spike-in” controls, which are three clonally derived gRNA plasmids that behave in known ways: a functional gRNA control, a nonfunctional gRNA control, and a wash control. The functional gRNA targets a site in *galM* that had been shown previously to be highly active^25^. Conversely the nonfunctional and wash controls contain a gRNA with no target in the genome, which have also been empirically tested. These controls are added to the donor library, or spiked-in, prior to transformation into the host, and thus they act as internal standards for expected behaviour within the library. The wash control, meanwhile, is spiked into the post-transformation recovery media to act as a proxy for contamination by plasmid DNA left over from transformation, which can and does interfere with PCR-based methods used for amplicon sequencing.

In conclusion, we speculate that the simple but validated design rules we developed here should be widely applicable for gRNA design because the sole metric used to judge the likely activity of a gRNA is derived from the spacer region. Because RNA tends to interact *in cis* and the spacer is on the 5’ end of the gRNA transcript, spacer self-interactions are formed long before the gRNA scaffold has disassociated from the RNA polymerase, let alone form interactions with Cas9 or the genome. Thus, spacer structure logically is a dominant feature contributing to gRNA function, as without RNA-DNA hybridization Cas9 cannot bind to the target DNA locus. This rule is also likely applicable to other nucleases, as they have similar requirements for their functionality though their differing spacer lengths mean different cutoff MFE values would likely have to be developed for each individual nuclease. Overall, this gRNA design approach represents a simple but empirical design strategy to avoid nonfunctional gRNAs in a practical, repeatable manner that could have applications across all organisms and highlights the large number of gRNAs that are functional in *E. coli*.

## ACKNOWLEDGEMENTS

Many thanks to Morgan Manning at Oak Ridge National Laboratory for his exceptional assistance with producing figures for this work.

## FUNDING ACKNOWLEDGEMENTS

This material is based upon work supported by the Center for Bioenergy Innovation (CBI), U.S. Department of Energy, Office of Science, Biological and Environmental Research Program under Award Number ERKP886. This work was also sponsored by the Laboratory Directed Research and Development Program of Oak Ridge National Laboratory, managed by UT-Battelle, LLC, for the US Department of Energy.

## DATA AVAILABILITY

Raw sequencing data generated in the course of this work has been deposited in GenBank under BioProject PRJNA1270398. Plasmid maps in GenBank format have been included in the Supplemental Material.

