## SupplementalMaterialAndData for "When good guides go bad: empirical evaluation of all unique Cas9 protospacers in *E. coli* reveal widespread functionality and rules for gRNA design": SupplementaryNote.docx

Supplementary Note – A region of the *E. coli* genome exhibits increased resistance to Cas9 activity

During the course of this work, we noticed the region between 280,000 and 290,000 bps on the *E. coli* chromosome was the sole region with an equal amount of functional and nonfunctional gRNA targets as well as an increase in the total number of unique gRNA targets. When we plotted the A-scores for each gRNA target by their genomic coordinates for the region between 265,000 and 305,000 bps from the origin, we observe a sharp decrease in gRNA functionality against targets in this 280,000 – 290,000 bp region compared to the surrounding genomic loci (Supplementary Figure 1A). Because of this highly localized region of nonfunctional gRNA targets, we nicknamed this region the Briar Patch. When we mapped the genomic features of the Briar Patch to the A-score plot, we found that the region is flanked by two identical insertion (IS) elements, specifically IS1B and IS1C. Hypothesizing that these flanking regions were playing some role in this depression of gRNA activity, we decided to 1) synthesize all the Briar Patch guides into a small library, 2) knock out IS1C with a selectable marker, then 3) transform our Briar Patch library into both the MG1655 wild-type host (Supplementary Figure 1B) side-by-side with our ISΔ host (Supplementary Figure 1C). When completed, we saw that the wild-type host recapitulated the depressed functionality we observed in the genome-wide library, while removing just one of the two IS elements relieved this depressed functionality.

While these results clearly indicate that the IS elements flanking the Briar Patch are responsible for this phenomenon, a couple of possible mechanisms might be responsible. The most obvious is composite transposition removing this region from the genome, though the size of the IS elements (768 bps) could be providing an escape route from Cas9 activity by homologous recombination repair at this region, which, while HR is an inefficient process in *E. coli* it does occur.

.
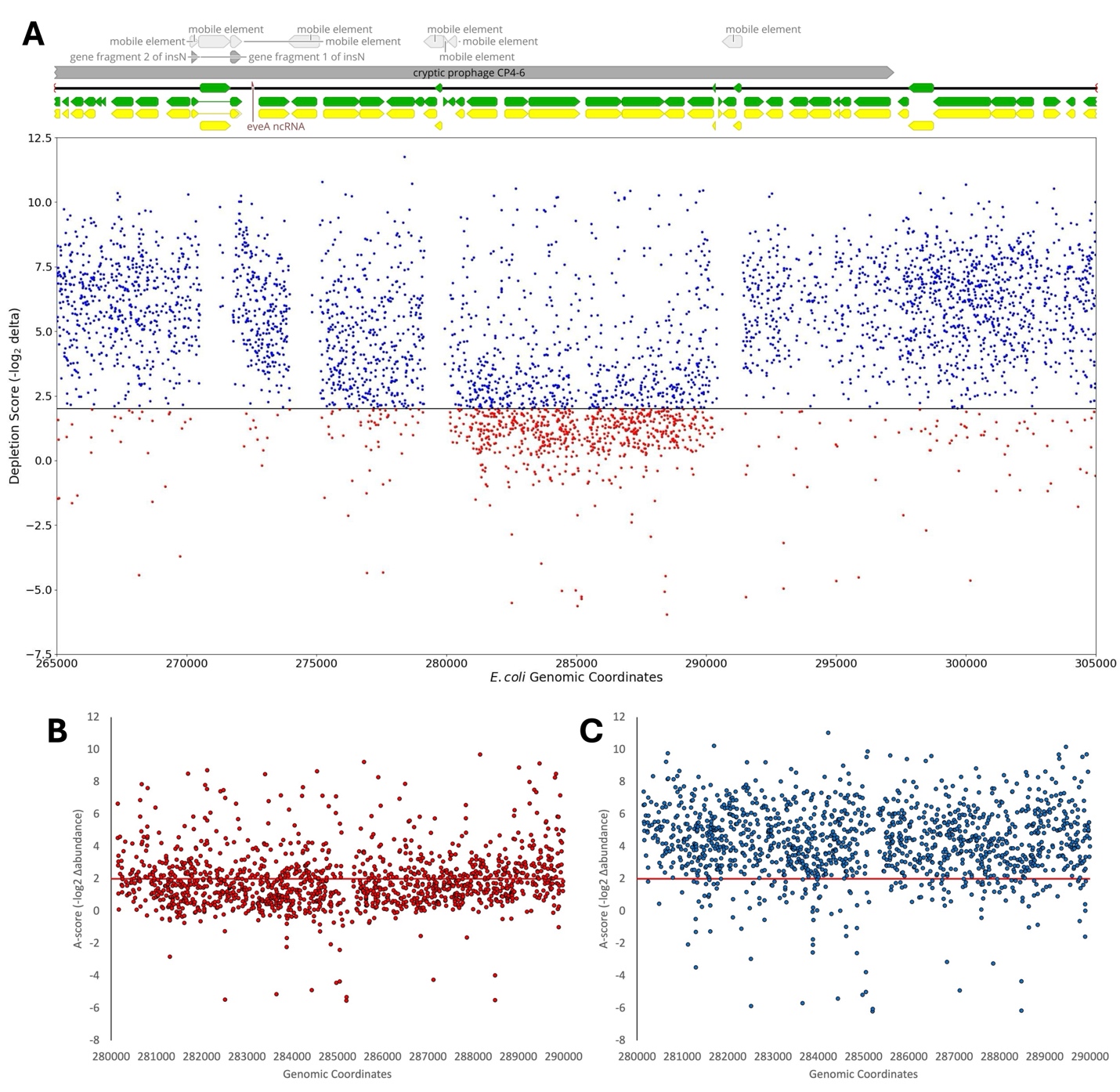

Supplementary Figure 1. The insertional elements flanking the Briar Patch are implicated in the apparent nonfunctionality of gRNAs targeting loci in this region. A) gRNA A-scores from the genome-wide library show a marked decrease in gRNA activity is observed between 280,000 and 290,000 bps. This boundary is precisely demarked by two identical *IS1C* elements, as seen by the genomic feature overlay at top. B) A-scores of Briar Patch library members transformed into the wild-type host. C) A-scores of Briar Patch library members transformed into the ISΔ host.
